# Investigating the mode of action for liver toxicity and wasting-like responses produced by high dose exposures to longer chain perfluoroacid substances (PFAS) using high throughput transcriptomics

**DOI:** 10.1101/2024.03.02.583129

**Authors:** A. Rasim Barutcu, Michael B. Black, Melvin E. Andersen

## Abstract

Single doses of perfluoro-n-decanoic acid (PFDA) cause wasting, a progressive loss of 30 to 50% body weight, increasing liver/body weight ratios, and death within several weeks (Olson and Andersen, 1983). Repeat high doses of perfluorooctane sulfonate (PFOS) produce a subset of these responses in rats and monkeys. The mode of action (MOA) of these wasting-like syndromes is not clear, nor is it understood if these responses are limited to a subset of perfluoroacid substances (PFAS) or a common response to high dose exposure with a larger number of PFAS. To identify pathway perturbations in liver caused by PFAS, we analyzed published *in vitro* gene expression studies from human primary liver spheroids treated with various PFAS for treatment times up to 14 days (Rowan-Carroll *et al*., 2021). With treatment times of 10 to 14 days, longer-chain PFAS compounds, specifically PFOS, perfluorodecane sulfonate (PFDS) and higher doses of perfluorooctanoic acid (PFOA), downregulated large numbers of genes in pathways for steroid metabolism, fatty acid metabolism and biological oxidations. Shorter chain PFAS compounds upregulated genes in pathways for fatty acid metabolism. Although PFDA was more toxic and could only be examined at 1-day of treatment, it also downregulated genes for lipid metabolism, steroid metabolism, and biological oxidations. Shorter chain PFAS, both carboxylic and sulfonic acids, did not lead to downregulation of pathways for fatty acid or steroid metabolism. TCDD is also known to cause wasting responses in rodents and humans. In intact rats, high dose responses of longer chain PFAS produce downregulation of batteries of genes associated with fatty acid oxidation and lipogenesis similar to those seen with TCDD. Based on our results, when combined with other literature, we propose that the longer-chain PFAS impair lipogenic pathways through inhibitory interactions between PPARβ, PPARα and PPARγ.

## Introduction

Wasting refers either to a syndrome where animals receiving a single dose of a substance progressively stop eating, lose weight, have increased liver:body weight ratios and die within several weeks of dosing or, on more prolonged exposure, have loss of body weight, anorexia and increased liver to body weight ratios. TCDD caused wasting in rodents (Gasiewicz *et al*., 1980; Christian *et al*., 1986) and anorexia and weight loss in workers exposed to TCDD during manufacture of trichlorophenoxyacetic acid (Poland *et al*., 1971; Girer *et al*., 2020). In a companion paper evaluating the mode of action for wasting seen with TCDD, female rats were treated with TCDD 5 days/week for 4 weeks, and the higher doses, 300 and 1000 ng/kg/day, caused wasting-like responses. Transcriptomic analyses of liver from these rats showed enrichment for downregulated genes in multiple pathways of cellular metabolism, including fatty acid, steroid, pyruvate metabolism and citric acid (TCA) cycle, and biological oxidations. The proposed mode of action for wasting caused by TCDD was extensive circadian cycle disruption (Andersen *et al*., 2024)

Perfluorodecanoic acid, PFDA, caused unequivocal signs of wasting in various rodent species after single doses of 50 to 100 mg/kg (Olson and Andersen, 1983; George and Andersen, 1986; Van Rafelghem *et al*., 1987). At these doses, rats had markedly reduced food intake, their body weight decreased by 30 to 40% over the first two weeks and they died 2 to 3 weeks after dosing. Compared to pair-fed controls, the liver to body weight ratios of the PFDA treated rats were over twice that in control animals on day 16 after dosing. Another longer chain PFAS, perfluorooctanesulfonic acid (PFOS) also caused wasting-like responses in rats and cynomolgus monkeys with longer term dosing. In 90-day sub-chronic toxicity studies, rats fed diets with 100 ppm or higher PFOS had liver enzyme elevations, hepatic vacuolization, hepatocellular hypertrophy, weight loss, emaciation, and died within 5 to 7 weeks (Goldenthal *et al*., 1978a; Seacat *et al*., 2003). When rats were dosed by gavage at 20 mg/kg/day for 4 weeks, there was weight loss, reduced food intake, and increased liver/body weight ratios (Cui *et al*., 2009). Similar cumulative toxicity with PFOS was observed in cynomolgus monkeys (Goldenthal *et al*., 1978b; Seacat *et al*., 2002), indicating that these wasting-like responses with PFOS, as noted with TCDD, occur in both rats and in primates. *In vitro* various PFAS activate both rat and human PPARα (a receptor involved in fatty acid oxidation) and PPARγ (involved in lipogenesis) (Evans *et al*., 2022) although it is unclear how activation of these pathways by persistent agonists would lead to wasting-like responses that have only been reported for longer chain PFAS and not for the shorter chain PFAS that also activate these same receptors.

The purpose of the present work was to take advantage of published studies of gene expression caused by various PFAS in human liver spheroids treated with various concentrations for up to 14-days (Reardon *et al*., 2021; Rowan-Carroll *et al*., 2021) and determine if longer chain PFAS caused similar changes in gene expression in liver tissue as those seen with TCDD in intact rats (Andersen *et al*., 2024). To caaccomplish this goal, we determined the dose response for differentially expressed genes (DEG) and evaluated Reactome pathway enrichment for up- and downregulated genes, looking for similarities in pathway enrichment between longer chain PFAS in these human hepatocyte liver cultures and those seen with TCDD in intact rats.

Our results reveal that treatment of the spheroids with longer chain PFAS caused significant enrichment with downregulated genes in pathways of lipid metabolism, fatty acid metabolism, steroid metabolism and biological oxidations, similar to the effects observed with TCDD (Andersen *et al*., 2024). We further integrated our results with other literature to offer a hypothesis for the mode of action (MOAs) for PFAS in causing wasting-like responses. Our hypothesis proposes that the longer chain PFAS, likely act through inhibitory actions of PPARβ,δ, causing transcriptional repression of both fatty acid metabolism and lipogenesis. Importantly, the downregulation of pathways for fatty acid metabolism, steroid metabolism and biological oxidations was not observed with shorter chain length PFAS where there was pathway enrichment with upregulated genes for fatty acid metabolism, biological oxidations, and regulation of lipid metabolism. There appears to be significant differences in biological modes of action for higher dose exposures of shorter (n < 7) and versus longer (n > 7) chain perfluorocarboxylic and perfluorosulfonic acids. Furthermore, while the wasting-like responses seen with TCDD and the longer chain PFAS appear to arise from profound downregulation of lipogenesis, the modes of biological action of these two classes of compounds likely differ.

## Materials and Methods

### Gene expression results with Human Primary Hepatocyte Spheroids

The altered gene expression in primary hepatocyte spheroids was from recent literature (Reardon *et al*., 2021; Rowan-Carroll *et al*., 2021). We analyzed the effect of chain length on DGE by examining perfluorocarboxylic acids (perfluoropentanoic acid (PFPenA), perfluorohexanoic acid (PFHxA), perfluoroheptanoic acid (PFHpA), perfluorooctanoic (PFOA), perfluorononanoic acid (PFNA), and perfluorodecanoic acid (PFDA) and several perfluorosulfonic acids - perfluorobutane sulfonic acid (PFBS), perfluorohexane sulfonic acid (PFHxS), perfluoroheptanesulfonic acid (PFHpS), PFOA and PFDS. In these studies, gene expression with the spheroids was measured using the human TempO-Seq S1500 panel from BioSpyder Technologies Inc, Carlsbad, California (House *et al*., 2017; Mav *et al*., 2018). The processed files were obtained from the NCBI Gene Expression Omnibus; series number GSE145239 for these various PFAS substances, and differentially expressed genes were determined by using the DeSeq2 package (Love *et al*., 2014). Pathway analysis was examined by using the Reactome ontology (https://reactome.org). We imported the Reactome pathway enrichment results into CytoScape 3.9.1 (Cytoscape.org) to generate GO-association network graphs, based on enrichment for up- and downregulated DEGs. In these GO-association network graphs, red nodes designate upregulated pathways, green nodes designate downregulated pathways and yellow represents nodes where there were both upregulated and downregulated genes in the pathway. In each graph, the sizes of the circles are proportional to the numbers of genes in the pathway.

Transcription factor binding site (TFBS) enrichment for DEGs used the 2022 ChEA database in Enrichr that allows reconstruction of a network of TFs based on shared overlapping targets and DNA-binding site proximity. The development and continuous enhancement of the Enrichr platform has been described in several publications (Chen *et al*., 2013; Kuleshov *et al*., 2016; Xie *et al*., 2021). The output from ChEA 2022 analysis was collected and used to create tables for TFs associated with different treatments for different times of exposure. The odds ratio used in the Enrichr plots is a statistical measure used to assess the enrichment of transcription factor binding sites (TFBSs) within the regulatory regions of genes in a specific gene set compared to a background reference gene set and indicates whether the TFBSs associated with a particular transcription factor is significantly overrepresented or underrepresented. Higher significance is associated with TFs with larger odds rations and lower *p*-values. The figures for CHEA 2022 analyses of DEGs plots both the odds ratio and p-value from the DEGs in the various Reactome categories.

### In-life studies with rats with PFOA

One aspect of the studies on wasting-like responses with PFAS is the questions of species differences with PFAS exposures. In a recent study (Gwinn *et al*., 2020), male Sprague-Dawley rats were treated for 5 days with PFOA at doses of 0.156, 0.325, 0.625, 1.25, 5, 10 and 20 mg/kg/day and gene expression determined using the TempO-Seq 1500 panel. We examined enrichment of DEGs at 5 and 20 mg PFOA/kg/day and evaluated pathway enrichment to compare results with PFOA seen in these rats *in vivo* with those seen with the primary human hepatocyte spheroids.

## Results

### Visualizing gene expression patterns caused by longer chain PFAS

Previous analysis of the DEGs of the gene expression results with thsee human liver spheroids identified pathway enrichment for cholesterol biosynthesis with PFOS and PFDS with downregulated genes and enrichment for fatty acid β-oxidation with upregulated genes for PFOA and PFBS (Rowan-Carroll *et al*., 2021). Possible *in vivo* MOAs were not discussed and pathway enrichment analyses for other PFAS were not reported. Using DEGs from their study, we examined the gene expression profile for 12 PFAS (Table 1) and evaluated the enrichment of various Reactome pathways for these DEGs.

**Table 1:**
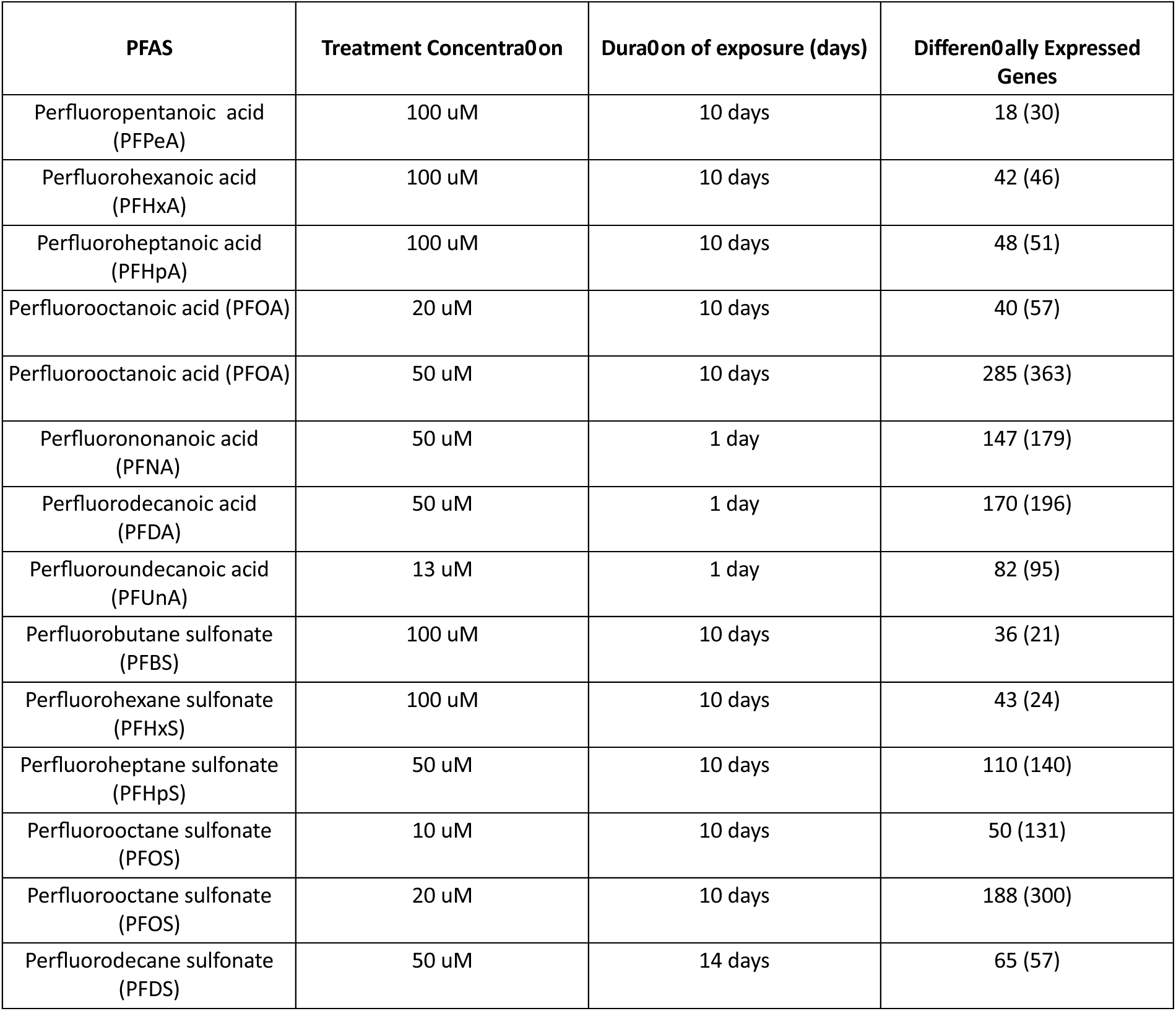
Data Sets used for assessing Pathway Enrichment for PFAS from published studies (Reardon *et al*., 2021; Rowan-Carroll *et al*., 2021) and the numbers of Up- and Down Regulated Genes for each PFAS/Treatment Regimen.

### Longer chain PFAS

With PFOS, we examined differential gene expression for 10 and 20 uM in spheroids treated for 10 days (**Figures 1 A-B**). At 10 uM there were 181 DEGs, and enrichment with downregulated genes for pathways of metabolism of steroids, cholesterol biosynthesis and biological oxidations. Among the genes downregulated in metabolism of steroids were genes coding for key proteins in the pathway from acetyl-CoA to lanosterol (*HMGCR, IDI1, FDFT* and *LSS*), genes for proteins from lanosterol to cholesterol (*CYP51A1, MSMOL* and *EBP*) and genes involved in cholesterol metabolism (*EBP, CYP7A1, CYP39A1, INSIG1,* and *HSD17B11*). (All affected genes in these pathways are listed in the figure legends.) At the 20 uM concentration, with 488 DEGs, there was more extensive enrichment of pathways, including downregulation of fatty acid metabolism and upregulation of multiple pathways consistent with increasing levels of cellular stress. Downregulated genes in the pathway for fatty acid metabolism include *ACADL* (long chain specific acyl-CoA dehydrogenase), *DECR1* (dienoyl CoA reductase), *EPHX2* (epoxide hydrolase 2), *ACYL* (ATP Citrate Lyase) and 7 *CYP* enzymes that participate in fatty acid metabolism. Two of these, *CYP4A11* and *CYP4A22*, catalyze the oxidation of fatty acids either through oxidation at the ω or ω-1 positions (Adas *et al*., 1999a; Adas *et al*., 1999b). *ACADL* is the first step in mitochondrial β-oxidation of long chain fatty acids; *DECR1* is involved in the mitochondrial oxidation of unsaturated fatty acids; *EPHX2* is involved in dephosphorylation of lyso-glycerophospholipids producing long chain fatty acids and *ACYL* is responsible for the synthesis of cytosolic acetyl CoA in many tissues. Cytosolic acetylCoA is used for synthesis of steroids, isoprenoids and fatty acids (Pietrocola *et al*., 2015; Shi and Tu, 2015). One of the upregulated daughter pathways for metabolism, metabolism of amino acids and their derivatives, was enriched with 23 upregulated genes. Of these 16 were genes coding for various ribosomal proteins, genes that were also represented in the upregulated node for metabolism of RNA.

**Figure 1:**
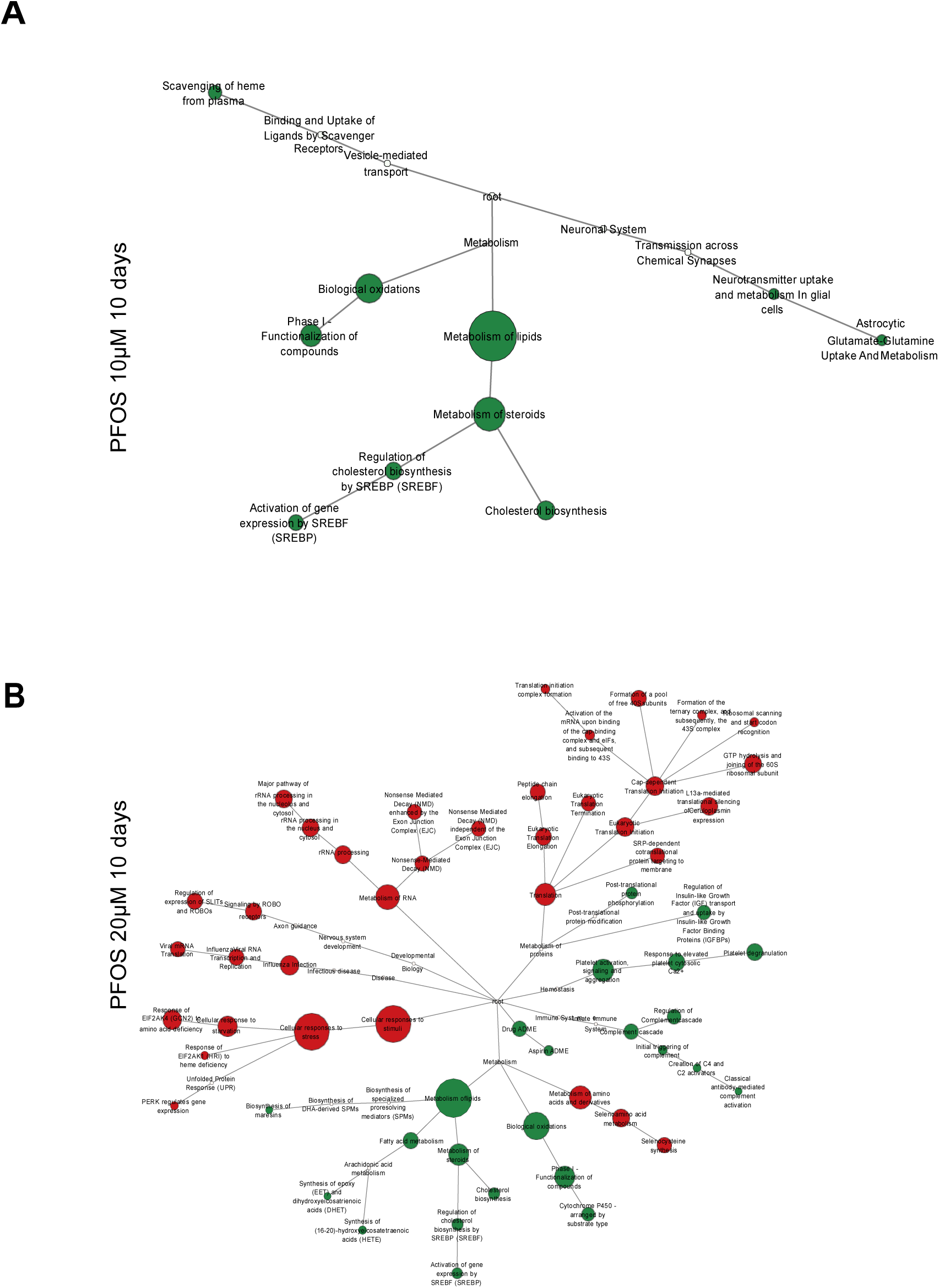
CytoScape visualization of pathway enrichment for genes altered by A) 10 uM PFOS for 10 days in human liver spheroids. Downregulated genes in metabolism of steroids pathway were: *ACAT2, ALB, CYP39A1, CYP51A1, CYP7A1, EBP, ELOVL6, FASN, FDFT1, HMGCR, HMGCS1, IDI1, INSIG1, LSS, MSMO1,* and *SLC10A1*. These genes regulate utilization of acetyl-CoA for cholesterol synthesis (ACAT2 and HMGCR), fatty acid synthesis (FASN and ELOVL6) and steroid hormone production. **B) Pathway map for 20 uM PFOS for 10 days in human liver spheroids.** Downregulated genes in fatty acid metabolism were: *ACADL, ACLY, CYP1A2, CYP2C19, CYP2C8, CYP2C9, CYP4A11, CYP4A22, CYP4F2, DECR1, ELOVL2, ELOVL6, EPHX2, FASN,* and *SCP2.* Some of these CYP enzymes are involved in the metabolism of long-chain fatty acids. In the metabolism of steroids pathway: *ACAT2, ALB, BAAT, CYP51A1, CYP7A1, DHCR7, EBP, ELOVL6, FASN, FDFT1, HMGCR, HMGCS1, HSD3B2, IDI1, INSIG1, LSS, MSMO1, NR1H4, SCP2, SLC10A1* and *SLCO1B1*.

With PFDS at a higher dose and longer time of treatment (14 days and 50 uM), there were 122 DEGs **(Figure 2)** with patterns of enrichment generally similar to that seen with PFOS except there was also downregulation of genes in the regulation of lipid metabolism by PPARα and no enrichment for upregulated genes in the pathway for cellular responses to stress. PFNA, PFDA and PFUnA were more toxic to the spheroids and DEG results were only reported for 1 day. Nonetheless, patterns of pathway enrichment with all three were similar to each other and to the longer-term treatments with PFOS and PFDS, showing downregulation of metabolism of steroids and biological oxidation (**Figure S1**).

**Figure 2:**
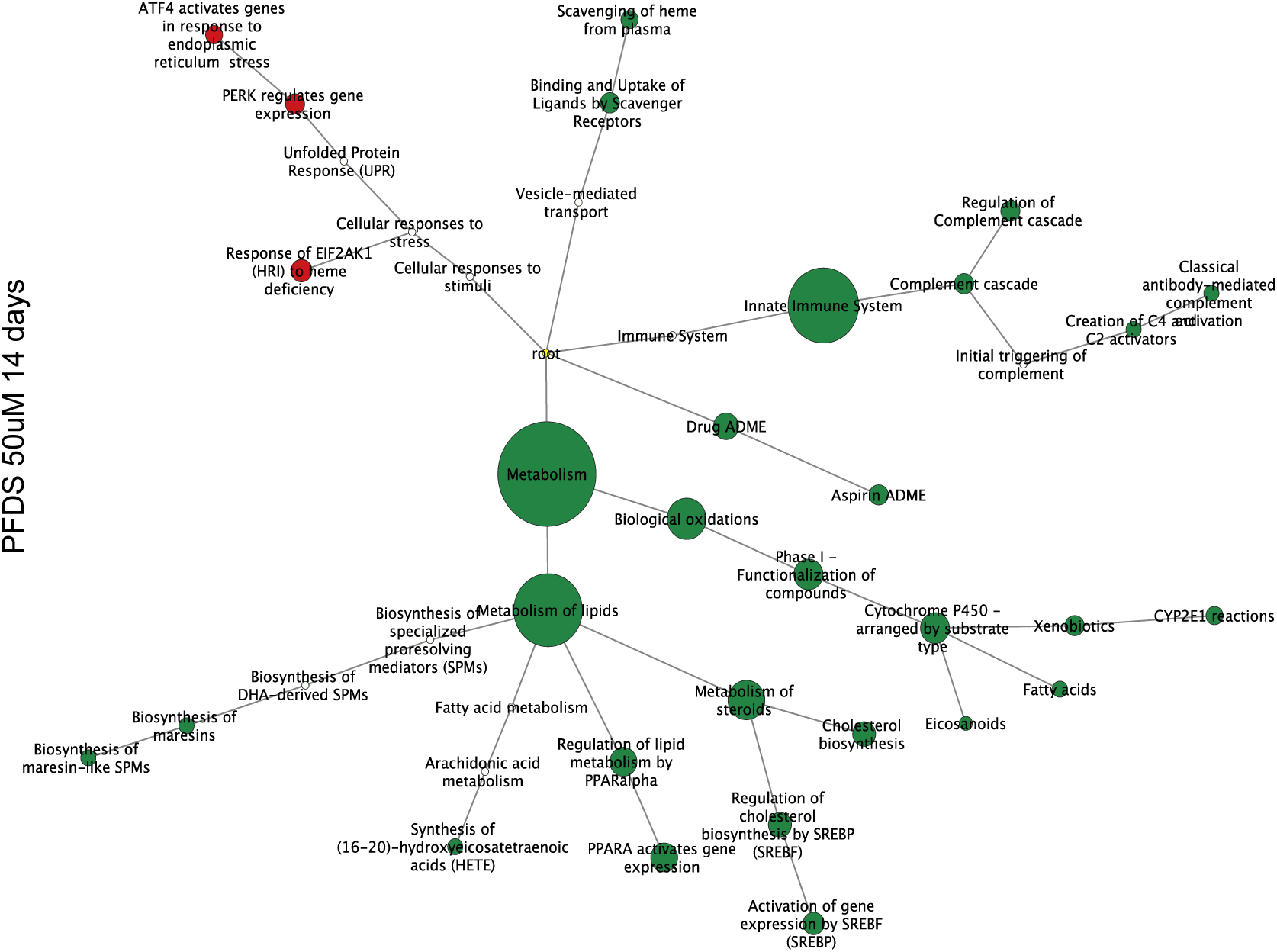
CytoScape visualization of pathway enrichment of genes altered by 50 uM PFDS for 14 days. Genes in metabolism of steroids pathway were *ACAT2, ALB, CYP51A1, EBP, ELOVL6, FASN, FDFT1, HMGCR, IDI1, INSIG1, LSS, MSMO1, CYP7A1, SLC10A1* and *LGMN*. In the regulation of lipid metabolism by PPARalpha pathway, *APOA1, FABP1, FDFT1, HMGCR, ME1, CYP7A1, HMGCS2, SULT2A1, ACADM* and *CYP4A11*were downregulated and *PLIN2, TNFRSF21, TXNRD1, TRIB3, PPARG, ACOX1* and *G0S2* were upregulated. In the Phase I-Functionalization of compounds, the downregulated genes were *CYP2C9, CYP2E1, CYP51A1, CYP2C8, CYP4F2, CYP7A1, CYP2A6, CYP2B6, CYP3A4, CYP4A11* and *CYP4A22*.

With PFOA at 10 days, there were qualitative differences as treatment concentrations increased from 20 to 50 uM. At 20 uM, there was enrichment for upregulated genes in pathways of fatty acid metabolism and PPARa activates gene expression pathways and pathway enrichment of metabolism of steroids with downregulated genes (**Figure 3A**). At 50 uM for downregulated genes, pathway enrichment was seen for metabolism of steroids, fatty acid metabolism and biological oxidations and in stress related pathways for upregulated genes (**Figure 3B**). The dose-dependent shift in enrichment for pathways of fatty acid metabolism was striking. At 20 uM, upregulated genes in the pathway included *ACADM, ACADVL, ACOX1, ACSL1, ACSL5, CYP4A11, CYP4A22, HADHA, PCCB* and *SLC27A2*. At 50 uM, there were 18 downregulated genes in this pathway. Of these only two, *CYP4A11* and *CYP4A22*, were upregulated at 20 uM. The downregulated genes included fatty acid synthetase (*FASN*) that synthesizes palmitic acid (C16:0) from acetyl and malonyl CoA; *ELOVL6* that synthesizes fatty acids with chain length greater than palmitic acid; and *ELVOL2* that participates in the synthesis of longer chain polyunsaturated fatty acids. Many of the downregulated CYP enzymes participate in hydroxylation of fatty acids. The gene expression pattern at 50 uM PFOA and with PFOS and PFDS indicates that the liver spheroids would have a greatly diminished capacity to synthesize a variety of lipids, steroids and isoprenoids. Among the downregulated genes at 50 uM was *SLC27A3*. These *SLC* genes code for fatty acid ligase proteins that conjugate free fatty acids with CoASH and facilitate transmembrane transport of the fatty acid esters. The other marked change in response going from 20 to 50 uM PFOA was enrichment for upregulated genes in various pathways, including cell response to stress and cell response to starvation.

**Figure 3:**
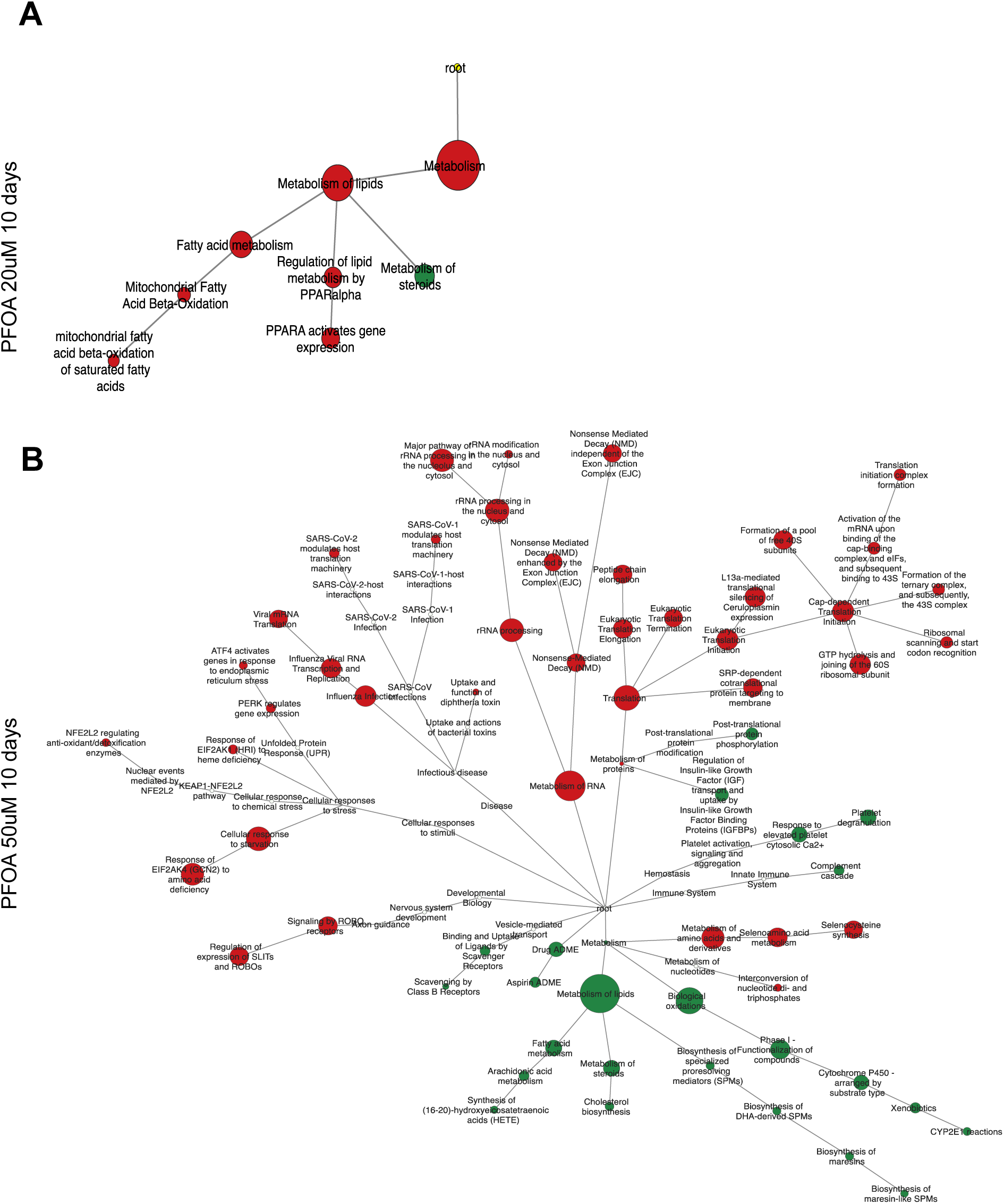
(A) CytoScape Visualization of altered Genes Expression in human liver spheroids for PFOA treatment at 20 uM for 10 days. Pathways for fatty acid metabolism and regulation of lipid metabolism by PPARalpha are enriched with enrichment for down regulated genes for steroid metabolism. Upregulated genes in the fatty acid metabolism pathway are: *ACADM, ACADVL, ACOX1, ACSL1, ACSL5, CYP4A11, CYP4A22, HADHA, PCCB,* and *SLC27A2.* Downregulated genes in the metabolism of steroids pathways are *ACAT2, CYP39A1, CYP51A1, CYP7A1, ELOVL6, HMGCR, IDI1,* and *SLC10A*. ELOVL6, fatty acid elongase enzyme 6, elongates fatty acids with 12, 14 or 16 carbons, with higher activity toward 16:0 fatty acids. (B) **Visualization of altered Genes Expression for PFOA treatment at 50 uM for 10 days.** The higher concentration of PFOA caused a major shift in gene expression. Genes in pathways for fatty acid metabolism and regulation of lipid metabolism by PPAR*α* are downregulated. Downregulated genes in fatty acid metabolism pathway are: *ACADL, ACLY, CYP1A2, CYP2C19, CYP2C8, CYP2C9, CYP4A11, CYP4A22, CYP4F2, DECR1, ELOVL2, ELOVL6, EPHX2, FASN, GGT1, GPX2, HADH, SCP2, SLC27A3*, and *HACD3*. Downregulated genes in the metabolism of steroids pathways are *ABCB11, ACAT2, ALB, CYP19A1, CYP39A1, CYP51A1, CYP7A1, EBP, ELOVL6, FASN, FDFT1, HMGCR, HSD17B11, IDI1, INSIG1, LSS, MSMO1, SCP2, SLC10A1* and *SLC51A*. Several genes in fatty acid metabolism were upregulated: *HADHA, ACADVL, ACSL1, ECI1, AKR1C3, THEM4,* and *ACSL5*.

A Venn diagram for downregulated genes for 20 uM PFDS, 50 uM PFOA and 20 uM PFOS (**Figure 4A**) showed 59 genes common to these three treatments. These downregulated genes were enriched for metabolism of steroids, metabolism of lipids, PPARA activates gene expression and biological oxidations. Downregulated genes for cytochrome P450 genes that code for proteins that metabolize fatty acids were *CYP2C8, CYP4F2, CYP2E1, CYP4A11,* and *CYP4A22*. We also performed an enrichment analysis for the genes unique to each of these compounds and to pairs of the three longer chain PFAS. For the individual compounds, interleukin-4 and interleukin-13 signaling and interferon gamma signaling had the largest numbers of genes/gene interactions with PFOS (102 genes); PPARa activates gene expression and cryoprotection by HMOX1 with PFDS (27 genes); and interleukin-4 and interleukin-13 signaling and neutrophil degranulation with PFOA (178 genes). For the treatment pair with high dose PFOA and PFOS (96 genes), there were increases in the number of genes/gene interactions for biological oxidations: *ADH1A, CAT, CYP2D6, FMO3, ADH1B, CYP1A2, CYP39A1, GSTM, BPHL, CYP2C19, FMO1* and *GSTM2*. Two of these *CYP*s*, CYP2D6* and *CYP2C19*, are involved in hydroxylation of long chain, polyunsaturated fatty acids (Lucas *et al*., 2010).

**Figure 4:**
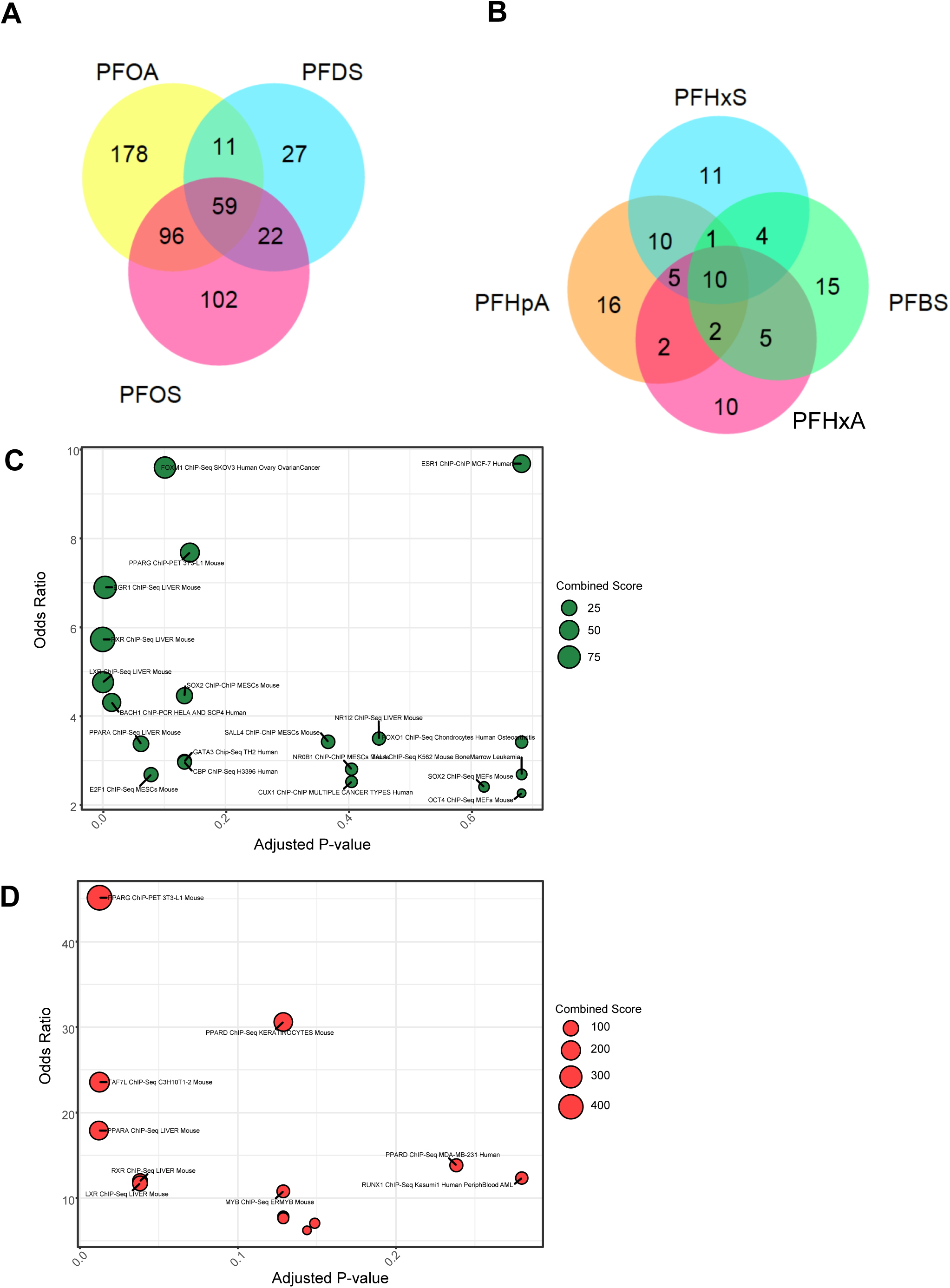
Venn diagrams for DEGs with shorter chain and longer chain PFAS: (A) Overlap of downregulated genes for PFDS (50 uM-14 days) and with the higher concentration treatments with PFOS (20 uM-10 days) and PFOA (50 uM-10 days) in human liver spheroids. With the downregulated 59 genes shared by all these compounds there was enrichment for gene expression by SREBP, metabolism of steroids, regulation of lipid metabolism by PPARalpha, and biological oxidations. B) Overlap for upregulated genes for PFHxA (100 uM–10 days), PFBS (100 uM – 10 days), PFHxS (100 uM-10days) and PFHpA (100 uM-10days). The 10 genes common to all in PPARA activates gene expression, were *ACADM, HMGCS2, ACSL1, PLIN2, CYP4A22, SLC27A2, POR, NRDG1, SGK2,* and *CYP4A11*. The genes unique to PFHxA for metabolism of steroids were *FASN, INSIG1, FDFT1, LSS* and *HMGCR*. (C) Enrichr plot for transcription factor binding sites for the 59 downregulated genes shared by PFOS, PFDS and the 50 uM PFOA groups. (D) Enrichr plot for the 10 up-regulated genes common to all groups in Panel B. The odds ratio is a statistical measure used to assess the enrichment of transcription factor binding sites (TFBSs) within the regulatory regions of genes in a specific gene set compared to a background reference gene set is plotted against another measure of significance, the-adjusted P-value.

Transcription factor binding site (TFBS) enrichment analysis of the 59 commonly downregulated genes (**Figure 4C**) found the most significant enrichment for RXR, LXR, PPARG and PPARA. RXR and LXR form heterodimers and act as nuclear receptors that regulate lipid metabolism, inflammation, and cholesterol homeostasis. Downregulation of these TFs indicates that treatment with higher doses of the longer-chain PFAS substances disrupt normal lipid homeostasis. Other enriched TFs, like FOXM1, SOX2, and E2F1 are involved in cell cycle regulation and proliferation.

Although some differences were observed in gene expression with these three longer chain PFAS, we found that at sufficiently high concentration in these human hepatocyte spheroids, all three produce consistent patterns of downregulation of genes in pathways for both steroid and fatty acid metabolism similar to a subset of transcriptional responses seen at higher doses of TCDD in rats – doses producing wasting-like responses (Andersen *et al*., 2024).

### Shorter chain carboxylic PFAS

With all shorter chain length carboxylic perfluorinated acids, responses of the spheroids were examined at 10 days. With PFPeA (data not shown), there were very few DEGs at 100 uM (18 upregulated and 30 downregulated) with enrichment for CYP2E1 reactions for upregulated genes and response to metal ions for downregulated genes. Curiously, 5 metallothionein genes were down regulated. With PFHxA (**Figure S3**), there was enrichment with upregulated genes for pathways of fatty acid metabolism, PPARα activates gene expression, and metabolism of steroids. The genes affected in the latter pathway were *ACAT2, FASN, IDI1, FDFT1, HMGCR, INSIG1, LSS, MSMO1* and *SLC27A2,* indicating activation of processes for both steroid synthesis and, through *FASN*, for fatty acid synthesis. With PHpA, (**Figure 5A**) there was upregulation of metabolism of lipids (20 genes) and PPARa activates gene expression with no enrichment of pathways for metabolism of steroids either with up or downregulated genes.

**Figure 5:**
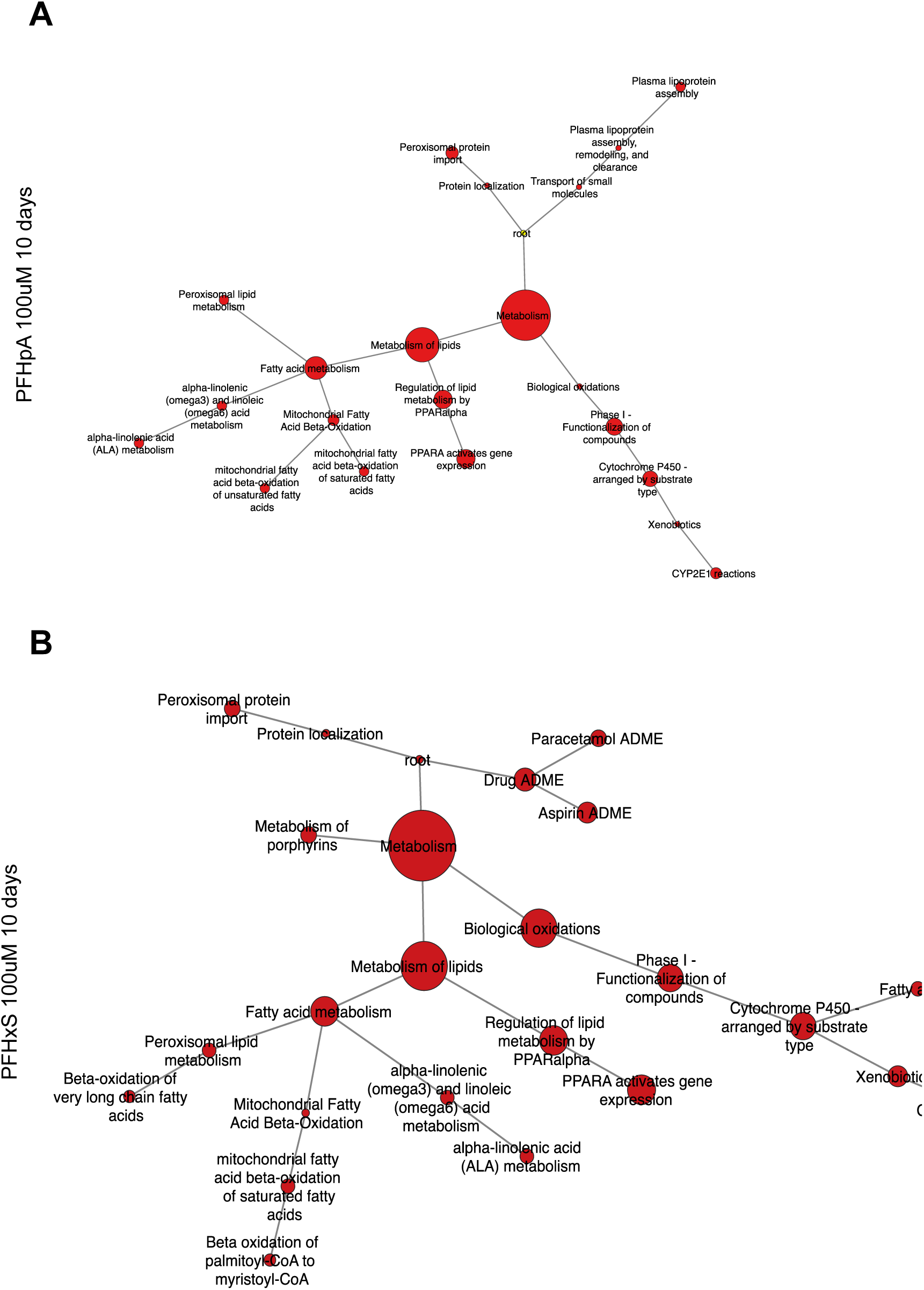
Visualizing Pathways enriched for differentially regulated genes for shorter chain Perfluorocarboxylic acids. A. **PFHpA**: Pathways for fatty acid metabolism and regulation of lipid metabolism by PPARalpha are enriched with no enrichment either in up or down regulated genes for steroid metabolism. The genes in the fatty acid metabolism pathway are: *ACAA1, ACADM, ACADVL, ACOX1, ACSL1, ACSL5, CYP2C19, CYP4A11, CYP4A22, DECR1, HADHA,* and *SLC27A2*. With PFHpA, acetylCoA is only being targeted for fatty acid metabolism through activation of fatty acid synthetase (FASN). B. **PFHxS at 100 uM for 10 days with human liver spheroids.** All pathways are enriched for upregulated genes and fatty acid metabolism upregulated genes were *ACAA1, ACADM, ACADVL, ACOX1, ACSL1, CYP2C8, CYP4A11, CYP4A22, HADHA,* and *SLC27A2*. Metabolism of steroid pathway genes were not affected with this PFAS. The pattern of altered gene expression with PFHxS resembles that of PFHpA; compounds of similar length (C_6_-O-SO_2_ and C_7_-CO_2_H).

**Figure 6:**
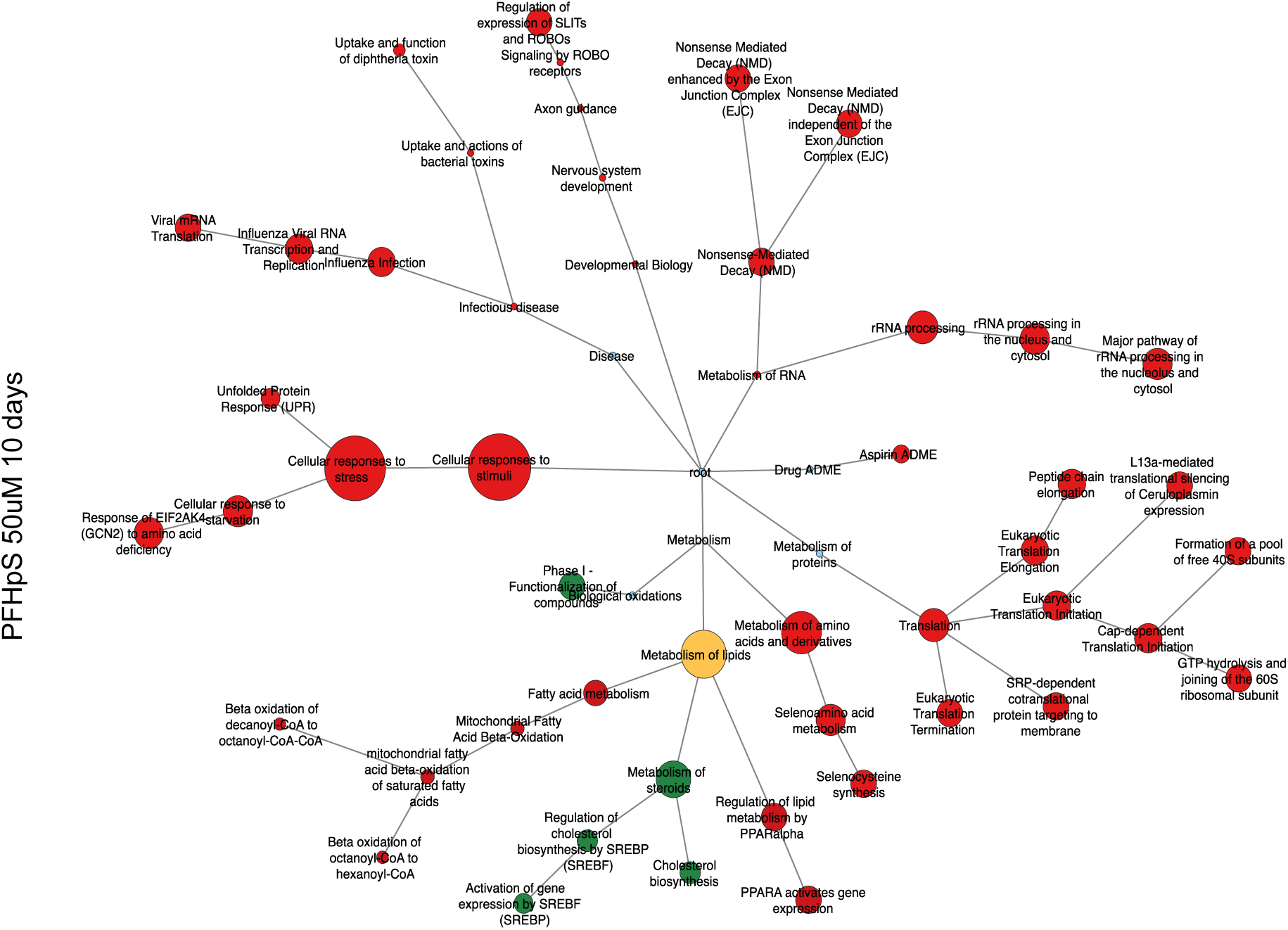
CytoScape visualization of pathway enrichment for DEGs with 50 uM PFHpS for 10 days. Fatty acid metabolism pathway upregulated genes were: *ACADM, ACADVL, ACOX1, ACSL1, ACSL5, AKR1C3, GPX2, HADH, HADHA,* and *SLC27A2*. The downregulated genes in the metabolism of steroid pathway were *ACAT2, ALB, CYP39A1, CYP51A1, CYP7A1, EBP, ELOVL6, FASN, FDFT1, HMGCR, IDI1, INSIG1, LGMN, LSS, MSMO1* and *SLC10A1*. This pattern with PFHpS has similarities to that with 50 uM PFOA with respect to the activation of cellular response to stress, metabolism of proteins, metabolism of RNA and infectious disease pathways.

### Shorter chain Sulfonic acid PFAS

A broadly similar pattern was evident with sulfonic acids. PFBS (**Figure S4**), which in terms of CF2 chain length is similar to PFPeA, showed enrichment in pathways for fatty acid metabolism, with 9 genes (including *ACAA1, ACADM ACSL1, SCP2* and *SLC27A2*) along with upregulation of 4 *CYP* genes. A similar pattern held with PFHxS (**Figure 5B**). With PFHpS (**Figure S5**), pathways for PPARa activates gene expression and fatty acid metabolism were enriched but not those for biological oxidations. At this chain length sulfonic acid, steroid metabolism was downregulated and, at the dose examined, there was upregulation of genes in pathways indicative of cellular responses to stress and toxicity.

In assessing similarities of DGE among upregulated genes for these 4 shorter chain length PFAS compounds, there were only 10 genes common to all of them and they are in the pathway for PPARa activates gene expression. For each, there was also upregulation of genes (**Figure 4**) not shared with other any other compound in the set. With the uniquely expressed genes with PFHxA (n=10), there was pathway enrichment for metabolism of steroids represented by seven genes: *ACAT2, HMGCR, MSMO1, FASN, INSIG1, FDFT1* and *LSS*, indicating likely increases in synthesis of cholesterol and long chain fatty acids. With PFHpA (n =16), there was pathway enrichment for lipoprotein clearance and metabolism of lipids, genes included in this pathway were *ACAT1, CYP2C19, ACSL5, DECR1, APOA1* and *G0S2*. ACAT1 catalyzes last step of the β-oxidation where it breaks down a fatty acid-CoA to acetyl-CoA and a fatty acid shorter by two carbons than the original substrate. ACSL5 converts long chain free fatty acids to their CoA-esters for either elongation or oxidation. DECR1 is a mitochondrial enzyme that participates in metabolism of unsaturated fatty acids in the mitochondria. The biological responses with PFHxA included upregulation of both fatty acid and steroid metabolism (PFHxA) while only fatty acid metabolism was affected with PFHpA. The picture is not as clear for the shorter-chain sulfonic acids. With PFBS, the pathway with the largest number of unique genes was metabolism with 9 genes: *BPHL, EPHX1, SCP2, CYP1A1, MAOA, SULT1B1, DDC, MDH1* and *TTR*. The pathway with the most genes uniquely associated with PFHxS was also metabolism with 7 genes: *AKR1C2, HSD17B11, UGT1A8, ALAS1, UGT1A10, GCLM* and *UGT1A4*.

Enrichment analysis of the commonly upregulated genes or with the 12 upregulated genes in fatty acid metabolism with PFHpA (data not shown), revealed significant enrichment for several transcription factors involved in lipid metabolism and adipogenesis, including PPARA, PPARG, PPARD, LXR and RXR. Their enrichment pattern indicates that perfluoroalkyl substance exposure activates genes are likely controlled by multi-factorial interactions with all three PPAR family members.

### Pathway Enrichment with PFOA in vivo in rats

Although PFOA has not been shown to cause wasting following treatment with a single oral dose, higher daily doses in rats or mice (Griffith and Long, 1980) led to decreased food consumption, decreased body weight and increased liver/body weight ratios. In a rat 90-day study at a dietary level of 1000 ppm, the mean liver/body weight ratios were 5.71 and 3.01 for the treated and control male rats. In male mice, the average liver to body-weight ratios in a 28-day study at 100 ppm in diet were 18.47 % and 4.61 %. With rats that survived until 14-days the liver to body weight ratios were 6.02 ± 0.28 (n=21) and 3.33 ± 0.08 (n=23)% for the 50 mg/kg and control groups respectively.

In short-term in life studies of differential gene expression with PFOA, male rats were dosed with 0.156, 0.325, 0.625, 1.25, 5, 10 and 20 mg/kg/day for 5 days (Gwinn *et al*., 2020). At the highest dose, there were 288 upregulated and 253 down-regulated genes with |FC|>1.5 and FDR<0.05 (Gwinn *et al*., 2020). As expected from past studies with compounds activating PPARα, there was enrichment for pathways of fatty acid metabolism and both mitochondrial fatty acid beta-oxidation and peroxisomal lipid metabolism for upregulated genes at both 2.5 and 20 mg/kg/day. At the higher dose, there was also pathway enrichment for cell cycle and mitotic G2-G2M phases with upregulated genes (pathways not shown in **Figure S6**). The enrichment for downregulated pathways included extracellular matrix and notably metabolism of steroids and cholesterol biosynthesis. There were 8 downregulated genes in the cholesterol biosynthesis pathway - *ACAT2, CYP51A, DHDCR7, HMGCR, IDI1, MSMO1, SQLE, TM7SF2* – and 11 in the metabolism of steroids, the eight above plus *SRD5A1, SCL10A2* and *CYP7A1*. Even in this short-term, in life study, there is clear evidence of downregulation of cholesterol synthesis pathway genes at daily doses where the total dose (100 mg/kg) is about half the single dose LD50 reported earlier (Olson and Andersen, 1983). There were increases in liver weight and liver to body weight ratios in the PFOA-groups of both 2.5 and 20 mg/kg/day and terminal body weights were significantly decreased for rats in the 20 mg/kg/day group. These *in vivo* DGE responses in male rat liver similar to the responses of human primary hepatocyte spheroids although the extensive enrichment of downregulated genes in pathways for metabolism of lipids as noted with the 50 uM treatment of spheroids (Fig. 3B) was not evident at the 20 mg/kg/day in vivo.

Overall, with the PFAS, either the carboxylic or sulfonic acids, there were similar trends (**Table 2**). The shorter chain length compounds show enrichment with upregulated genes for pathways of fatty acid metabolism, biological oxidations, and PPARa activates gene expression (though there were differences among the group as noted in **Figure 4**. With some combination of increasing chain length and exposure concentration, there was a shift where fatty acid metabolism and PPARa activates gene expression are upregulated and metabolism of steroids becomes downregulated (PFOA 20uM and PFHpS 50 uM). With the higher concentrations of PFOA and with both PFOS and PFDS, pathways for fatty acid metabolism, biological oxidations and metabolism of steroids are enriched with downregulated genes and pathways for cell response to stimuli and cell response to stress.

**Table 2:**
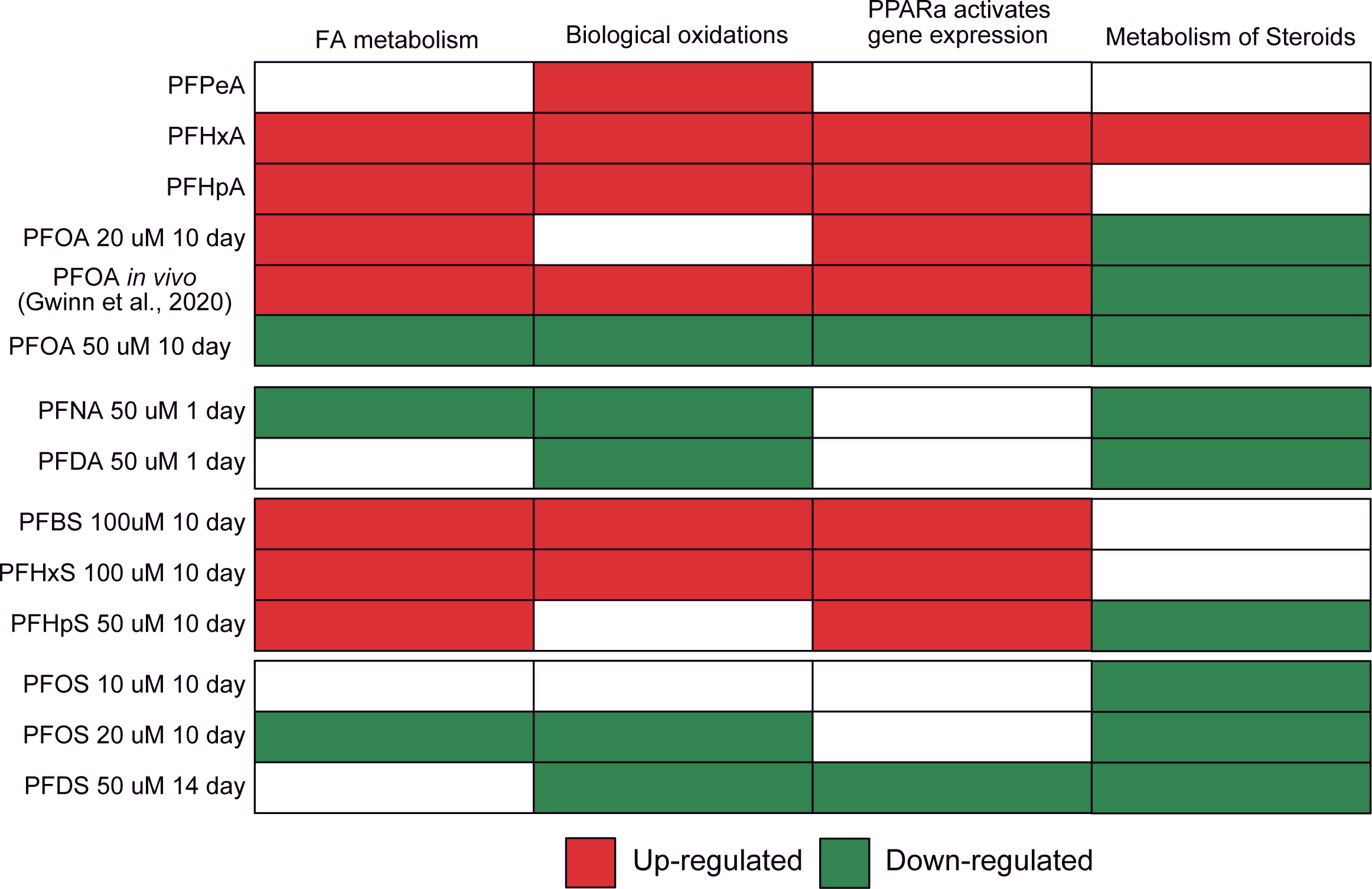
Summary of Regulation of most significanly enriched pathways of lipid metabolism by PFAS. Red indicates upregulation of pathway genes; green is downregulation.

Another gene downregulated with the longer chain PFAS was *SLC13A5*, a sodium dependent transporter that mediates transport of citrate into hepatocytes (Gopal *et al*., 2007). Citrate functions in various pathways through actions of ATP-citrate lyase (ACYL) that facilitates synthesis of acetyl-CoA for use in fatty acid metabolism, steroid synthesis, energy production, and gluconeogenesis. *ACYL* was downregulated with PFOA (50 uM), PFHpS, and PFOS 20 uM. This pattern of pathway enrichment for downregulated genes for lipogenesis and fatty acid metabolism with these longer chain PFAS closely aligns with high dose effects of TCDD on these pathways. Not unexpectedly, there was TFBS enrichment for PPARA and PPARD for many of the differentially expressed upregulated genes with the shorter chain length PFAS **(Figure 4D)**. The Enrichr analysis with the commonly downregulated genes in Figure 4B showed TFBS enrichment for PPARG, PPARA and PPARD.

The chain-length dependent shift in responses, from upregulation of genes for fatty acid metabolism to down regulation of genes for steroid metabolism, occurred between 20 and 50 uM concentrations with PFOA and with between PFHpS and PFOS (Figure 3 and Figure S5). Comparing Enrichr plots for TFBS with the upregulated genes in metabolism of lipids with next shorter pairs - PFHpA and PFHxS (Fig S2) - showed more significant enrichment for PPARD than was evident examining the common upregulated genes for the 4 shorter chain PFAS (Figure 4D).

## DISCUSSION

### Metabolic Targets for Wasting with longer-chain PFAS

Higher doses of PFOS, caused weight loss and liver enlargement in vivo in rats and non-human primates. The gene expression changes in the human liver spheroids showed pathway enrichment for downregulated genes in metabolism of fatty acids, metabolism of steroids, and biological oxidations in human liver spheroids. In comparison, TCDD affected these same pathways, but affected a much larger number of genes in these pathways (Andersen *et al*., 2024). While some of these differences in numbers of DEG affected may arise from the different platforms – full genome Affymetrix chips with TCDD versus TempO-Seq - TCDD also affected a much larger set of pathways than did PFOS. Another difference between TCDD and PFOS was changes in liver cell proliferation. With TCDD at 300 ng/kg/day, Ki67 staining revealed nearly complete cessation of cell proliferation (Andersen *et al*., 2024). With PFOS, no decreases in proliferating cell nuclear antigen (PCNA) staining had been noted in either rats or cynomolgus monkeys (Seacat *et al*., 2002; Seacat *et al*., 2003). The differential effects on cell proliferation are the likely explanation for the much larger increases in liver weight with PFAS (little effect of proliferation) and TCDD (almost complete inhibition of cell proliferation in liver).

### Altered Gene Expression and Wasting

The test systems used for evaluating gene expression of TCDD in rats and PFAS in human hepatocyte spheroids vary considerably, including analysis of different species (human versus rat), in-life with TCDD in the rats versus *in vitro* exposures with the human liver spheroids, and a unicellular platform versus exposure in an intact animal where the outcome will be affected by the multicellular milieu of the whole liver. Nonetheless, the high dose DEG enrichment patterns for TCDD and for the longer chain PFAS do share key changes in pathway enrichment with downregulated pathways for metabolism of lipids, fatty acid metabolism, metabolism of steroids, and biological oxidations. With TCDD, the lower doses (22 ng/kg/day in CL and 100 ng/kg/day in PP) upregulated genes were primarily controlled by AhR and Arnt – among these were several Cyp proteins (*1a1, 1a2* and *1b1*) and *Nqo1*- without downregulation of pathways of cellar metabolism. Only at 300 and 1000 ng/kg/day was there evidence for upregulation of pathways associated with cellular stress and down regulation of multiple pathways of cellular metabolism. Enrichr results plots with the higher doses of TCDD showed statistically significant contributions of PPARA, LXR and RXR without the signals for PPARG seen with the longer chain PFAS.

A strong case can be made that wasting and wasting-like syndromes, at least at the level of altered metabolic processes, with TCDD and with the longer chain PFAS, is associated with profound downregulation of lipogenesis and fatty acid metabolism. The observed enrichment of fewer pathways with longer-chain PFAS than with TCDD indicates that disruption of a more targeted group of pathways affecting lipogenesis may be sufficient to cause wasting-like responses. The proposed MOA for wasting with TCDD was based on interruption of circadian cycling by persistent upregulation of AhR-Arnt regulated genes and subsequent failure to move from the activity to the rest phases of the circadian cycle (Andersen *et al*., 2024). With longer-chain PFAS, the MOA for wasting-like responses appears to relate to inhibition of a more limited set of lipogenic pathways.

### MOAs for Effects of PFAS on Gene Expression and Wasting

Nearly 40 years have now passed since the first descriptions of wasting by PFDA (Olson and Andersen, 1983; George and Andersen, 1986). These 40 years have seen remarkable advances in understanding the role of the PPAR family of receptors (PPARα, PPARγ and PPARβ,γ) play in the biological responses to administration of these compounds and in the tools of biological research such as gene expression analyses. The present studies with the longer chain PFAS support the contention that prolonged inhibition of lipogenesis causes wasting and wasting-like responses. They do not, however, provide a clear understanding of the biology that is affected by these compounds, leading to downregulation of these pathways. We believe that our results of altered gene expression, when integrated with an emerging understanding of the function of these receptors, provides a plausible hypothesis for the mode of action of the longer chain PFAS in causing wasting, emphasizing that this is a hypothesis, not proven conjecture, that is consistent with known biology and suggestive of avenues of further research.

### Hypotheses for molecular targets leading to altered lipid metabolism and wasting (PFAS)

With longer chain length PFAS there was pathway enrichment with downregulated genes for metabolism of steroids, fatty acid metabolism and biological oxidations (**Table 2**). This pattern would be unexpected if responses of these PFAS were due solely to activation of either PPARα (upregulation of fatty acid oxidation) or PPARγ (upregulation of lipogenesis). The Enrichr analysis of TFBS with both upregulated genes for the shorter-chain PFAS and downregulated genes with the longer chain PFAS indicated contributions from multiple PPAR family proteins. The function of the third transcription factor in the PPAR family, PPARβ,δ, is not as well-established although this TF has activity as a transcriptional repressor and has been suggested to be an integrator of PPAR family receptor signaling (Shi *et al*., 2002). Studies with PPARδ−KO mice suggested that PPARδ regulates circadian changes in hepatic lipogenesis (Wang *et al*., 2020). On its own, PPARδ repressed basal transcription and, when co-expressed with PPARα or PPARγ, led to inhibition of specific target genes for these two receptors, showing both inhibition of genes coding for enzymes involved in fatty acid oxidation and lipogenesis (Shi *et al*., 2002). The inhibition of activation by either PPARα or PPARγ was due to the ability of unliganded PPARδ to recruit various corepressors affecting histone acetylation. The ability of liganded PPARδ to modulate PPARα and PPARγ gene expression has not been examined.

Using a variety of synthetic ligands (Wang *et al*., 2020), the ligand binding pocket for PPARδ, was shown to have three regions. The core portion of the binding site was optimally activated with a hexanoate group bound through a phenolic linkage to a substituted benzene moiety. Shorter chain lengths showed lower activity. The hexanoate head group and two carbons in the phenolic anchor would be equivalent to a C-8 or 9 chain length perfluorinated carboxylic side chain or a 7 or 8 chain length perfluorinated sulfonic acid side chain. These results are consistent with a hypothesis that the downregulation of metabolism of steroids and fatty acid metabolism by binding of these longer chain PFAS to PPARδ and action of the liganded PPARδ as a transcriptional repressor of both PPARα and PPARγ (Shi et al, 2002). However, saturated FAs, through lauric acid (i.e., dodecanoic acid) were reported to have binding constants >30 uM using a scintillation proximity assay with PPARα, PPARγ, and PPARδ (Xu *et al*., 1999). No mention was made of whether even shorter chain saturated fatty acids were tested, and the gene expression results here indicate that genes regulated by PPARα and PPARγ are increased by various shorter chain PFAS and then downregulated with increasing chain length or, in the case of PFOA or PFHpS, downregulated with increasing treatment concentration. Coupled with the observation that the genes downregulated by the longer chain PFAS have TFBS for both PPARA and PPARG support more complex interactions than simply effects on a single PPAR complex alone. We propose that this downregulation likely occurs due to inhibitory interactions of PPARδ on both PPARα and PPARγ, a response that depends on PFAS-chain length.

Some toxic responses to PFOA are PPARα independent. Slight delays in eye opening with PFOS were similar in PPARα-WT and PAPRA-KO mice suggesting that PPARα activation was not necessary for this effect (Abbott *et al*., 2009). Similarly, low dose gestational exposures to PFOA led to PPARα independent liver effects as the mice aged (Filgo *et al*., 2015). Liver gene expression due to various PFAS was evaluated in both WT and PPAR-KO mice (Rosen *et al*., 2017) and 11 to 24% of the DEGs were independent of PPARα. These PPARα independent results should not be unexpected for longer chain PFAS where there were significant numbers of downregulated pathways and genes in the human liver spheroids and the more varied regulation of these genes suggested by our EnrichR analysis of TFBS.

Different phases of the circadian cycle require diurnal adjustments of metabolism. In the activity phase a key pathway for production of energy is fatty acid oxidation leading to acetylCoA that can be used in the Krebs cycle for synthesis of ATP. During the resting portion on of the cycle, there is a need to synthesize longer chain fatty acids to produce structural lipids to support tissue repair, cell proliferation and growth. The synthesis of these lipids begins with acetylCoA (Pietrocola *et al*., 2015). The synthesis of longer chain fatty acids takes place in the cytoplasm and the synthesis of longer chain fatty acids occur with a growing chain complexed by acyl carrier protein. Very small amounts of free fatty acids above C4 are present in the cell and FASN holds the growing fatty acid eventually producing palmitate (C16). The PFAS studied here are structurally similar to fatty acids. There is little evidence that these PFAS are metabolized (Vanden Heuvel *et al*., 1991a; Vanden Heuvel *et al*., 1991b; Goecke *et al*., 1992; Kemper and Nabb, 2005). These compounds could be carried across cell and organellar membranes and interfere with β-oxidation or serve as a signal to enhance the oxidation and decrease free fatty acid concentrations. It is more difficult to see how longer chain PFAS signal that the cell has an adequate or excess of various lipids and generate feedback to diminish pathways of fatty acid and steroid synthesis. Perhaps, the longer chain fatty acids serve as negative regulators of metabolism by mimicking native regulatory fatty acids. In formulating risk assessment approaches for PFAS a key challenge will be gaining a better understanding of the underlying biology of processes that serve as control points for fatty acid and steroid metabolism and how they are affected by PFAS exposures. Lastly, with respect to MOAs of wasting in rats, subsequent studies should examine gene expression in rats dosed with PFDA at doses causing wasting to determine the similarities in DEGs with TCDD. In addition, future reseach could examine protein levels of key proteins where there was alteration in gene expression and utilize transgenic animals lacking *PPARδ*.

## Acknowledgements

The detailed analysis of the gene expression for PFAS compounds was supported by a grant from the Foundation for Chemistry Research and Initiatives, a 501(c)(3) organization established by the American Chemistry Council (ACC). The Long-Range Research Initiative of the American Chemistry Council (ACC-LRI) has provided support to Hamner Institutes and now ScitoVation over a longer period for developing and applying tools for assessing modes of actions for chemicals based on gene expression profiling.

## Supplemental Figure

**Figure S1: CytoScape Visualization for pathway enrichment of DEGs from human liver spheroids treated with A) PFNA and B) PFDA at 50 uM for 1 day and C) PFUnA at 13 uM for 1 day.**

With these compounds, the only treatment time without toxicity was that for 1-day. Downregulated genes in the metabolism of steroids pathways for PFNA were: ACAT2, CYP39A1, CYP51A1, CYP7A1, EBP, FASN, FDFT1, HMGCR, HSD17B11, HSD3B1, HSD3B2, IDI1, INSIG1, MSMO1, SCP2, and SLC10A1. With PFDA they were: ACAT2, CYP39A1, CYP51A1, CYP7A1, EBP, FASN, HMGCR, HSD3B1, HSD3B2, IDI1, INSIG1, LSS, MSMO1, NSDHL, SCP2, SLC10A1, and SLCO1B1. With PFUnA, they were EBP, CYP39A1, HMGCR, MSMO1, CYP7A1, FASN, CYP51A1, LSS, FDFT1, ACAT2, IDI1, SCP2, INSIG1, and SLC10A1.

**Figure S2: Enrichr Analysis of TFs enrichment for A) PFHpA with 14 genes in the metabolism of lipids pathway and B) for PFHxS with 19 genes in the metabolism of lipids pathway.**

The odds ratio, a statistical measure used to assess the enrichment of transcription factor binding sites (TFBSs) within the regulatory regions of genes in a specific gene set compared to a background reference is plotted against another measure of significance, the adjusted P-value.

**Figure S3: CytoScape Visualization for pathway enrichment of DEGs from human liver spheroids treated with PFHxA at 100 uM for 10 days.**

Pathways for fatty acid metabolism and metabolism of steroids are enriched for upregulated genes. The upregulated genes in metabolism of steroids included: ACAT2, FASN, FDFT1, HMGCR, IDI1, INSIG1, LSS, MSMO1, and SLC27A2. With the fatty acid metabolism pathway, the genes were: ACADM, ACADVL, ACOX1, ACSL1, CYP2C9, CYP4A11, CYP4A22, CYP4F2, FASN, HADHA, HSD17B8, and SLC27A2. With this compound, acetylCoA utilizing pathways are activated for synthesis of isoprenoids, cholesterol, and fatty acids.

**Figure S4: CytoScape Visualization for pathway enrichment of DEGs from human liver spheroids treated with PFBS at 100 mM for 10 days.**

The 27 upregulated genes in metabolism were ACAA1, ACADM, ACSL1, ALDH2, BPHL, CYP1A1, CYP2C8, CYP2E1, CYP3A4, CYP3A5, CYP4A11, CYP4A22, DDC, EPHX1, GLUL, HMGCS2, IDI1, MAOA, MDH1, PLIN2, POR, SCP2, SLC27A2, SULT1B1, SULT2A1, TST, and TTR. Nine of these, ACAA1, ACADM, ACSL1, CYP1A1, CYP2C8, CYP4A11, CYP4A22, SCP2, and SLC27A2 were in fatty acid metabolism pathway.

**Figure S5: CytoScape Visualization for pathway enrichment of DEGs from human liver spheroids treated with 50 uM PFHpS for 10 days.**

Fatty acid metabolism pathway upregulated genes were: ACADM, ACADVL, ACOX1, ACSL1, ACSL5, AKR1C3, GPX2, HADH, HADHA, and SLC27A2. The downregulated genes in the metabolism of steroid pathway were ACAT2, ALB, CYP39A1, CYP51A1, CYP7A1, EBP, ELOVL6, FASN, FDFT1, HMGCR, IDI1, INSIG1, LGMN, LSS, MSMO1 and SLC10A1. This pattern with PFHpS has similarities to 50 uM PFOA with respect to the activation of cellular response to stress, metabolism of proteins, metabolism of RNA and infectious disease pathways.

**Figure S6: CytoScape Visualization for pathway enrichment of DEGs from an in-life study in male Rats dosed with 20 mg/kg/day for 5 days (Gwinn *et al*., 2020).**

Downregulated genes in “metabolism of steroids” were: Hmgcr, Tm7sf2, Dhcr7, Cyp7a1, Srd5a1, Idi1, Cyp51, Acat2, Msmo1, Slc10a2, and Sqle. There were 23 upregulated genes in the fatty acid metabolism pathway: Acot1, Elovl6, Cyp2j4, Crat, Cpt1b, Acot4, Hadhb, Cpt2, Decr1, Prkab2, Acadvl, Slc27a2, Cyp4a1, Acaa2, Eci2, Acadm, Ehhadh, Acadl, Hacl1, Crot, Cbr1, Acox1, and Hadh. This plot is a smaller subset of pathways altered by PFOA which also induced cell proliferation at these higher doses. The in-life analysis for gene expression with PFOA was done with the Temp-O-seq platform.

