## Supplementary figures and images for "Investigating the mode of action for liver toxicity and wasting-like responses produced by high dose exposures to longer chain perfluoroacid substances (PFAS) using high throughput transcriptomics"

### Supplemental Materials

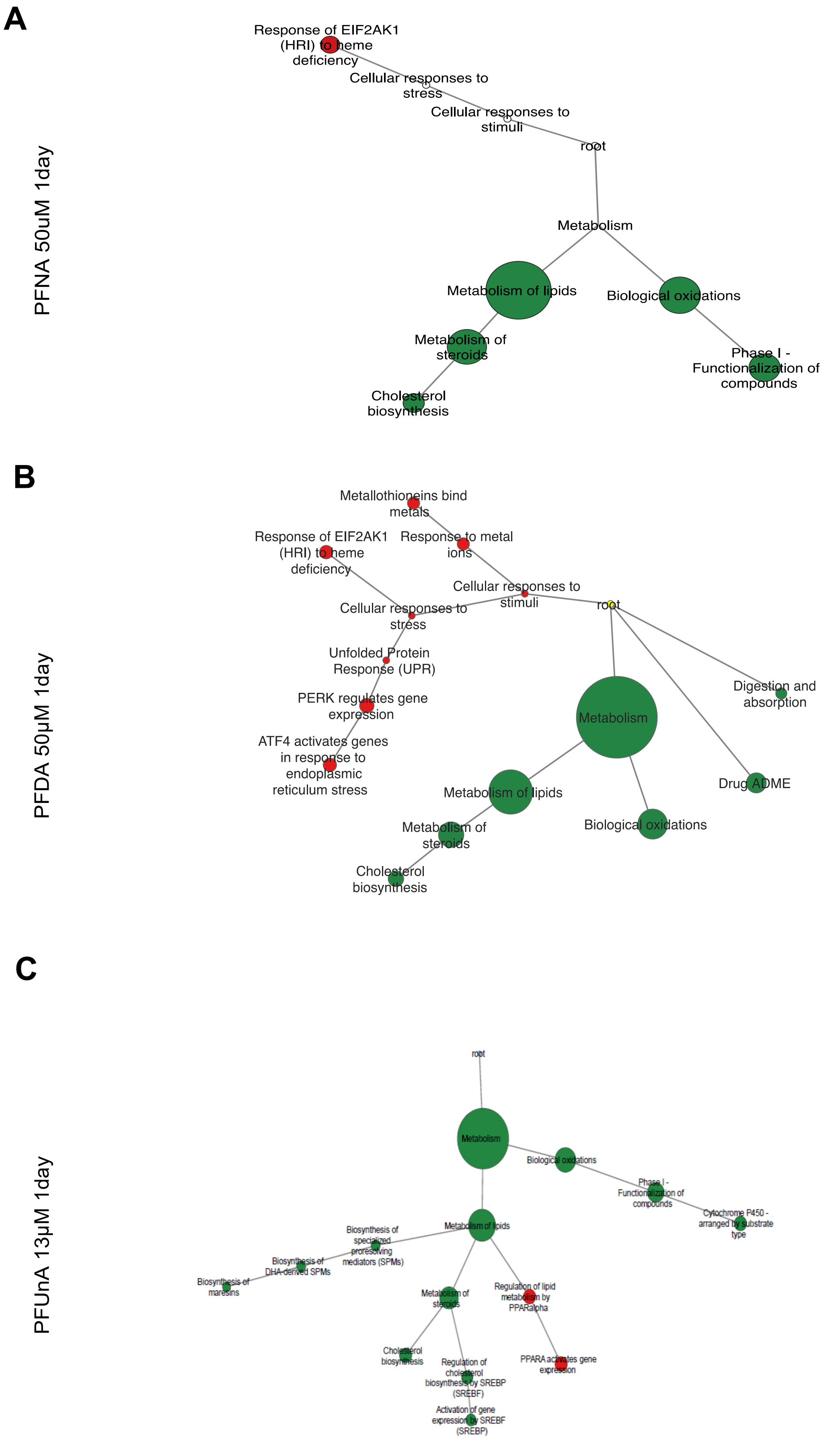

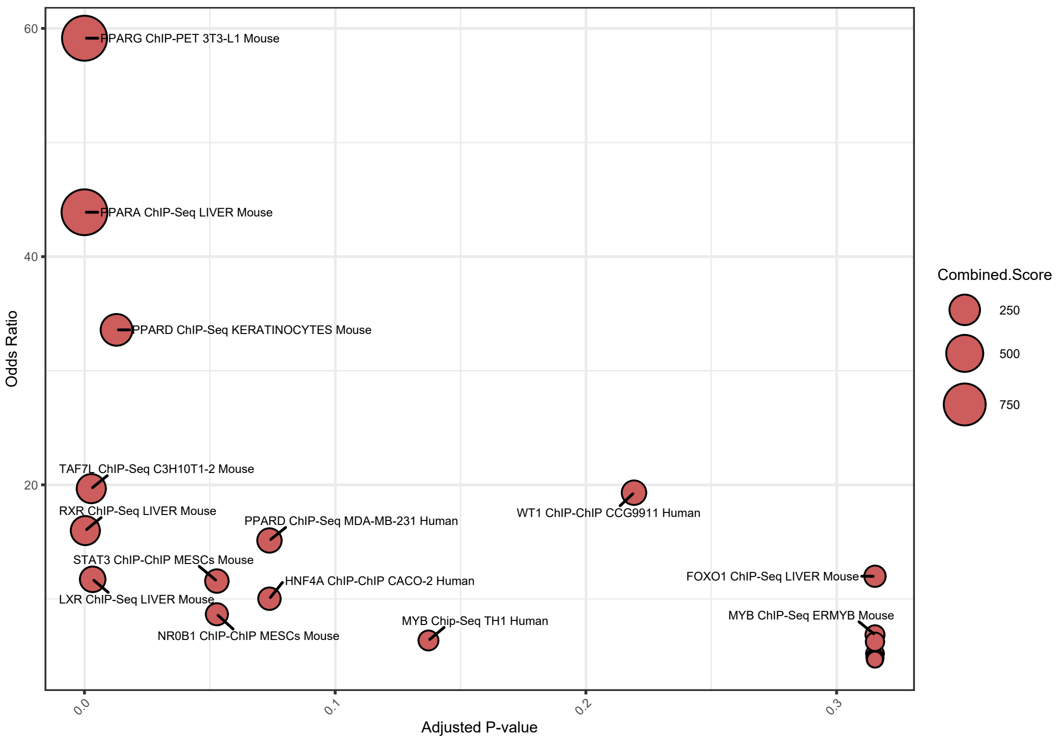

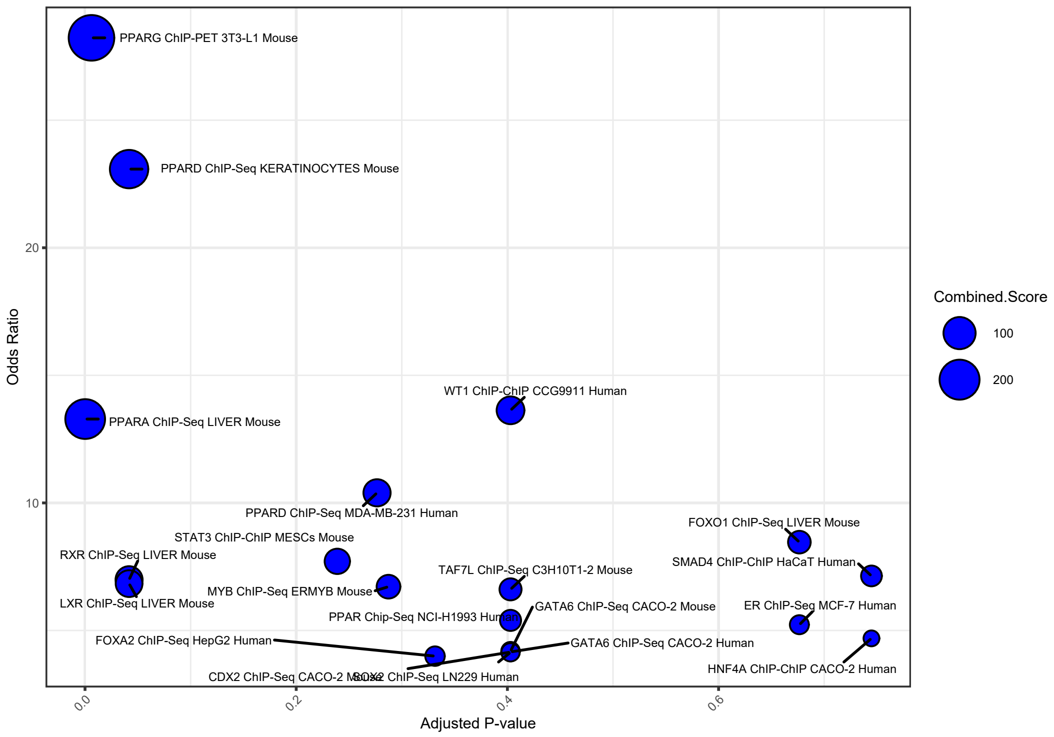

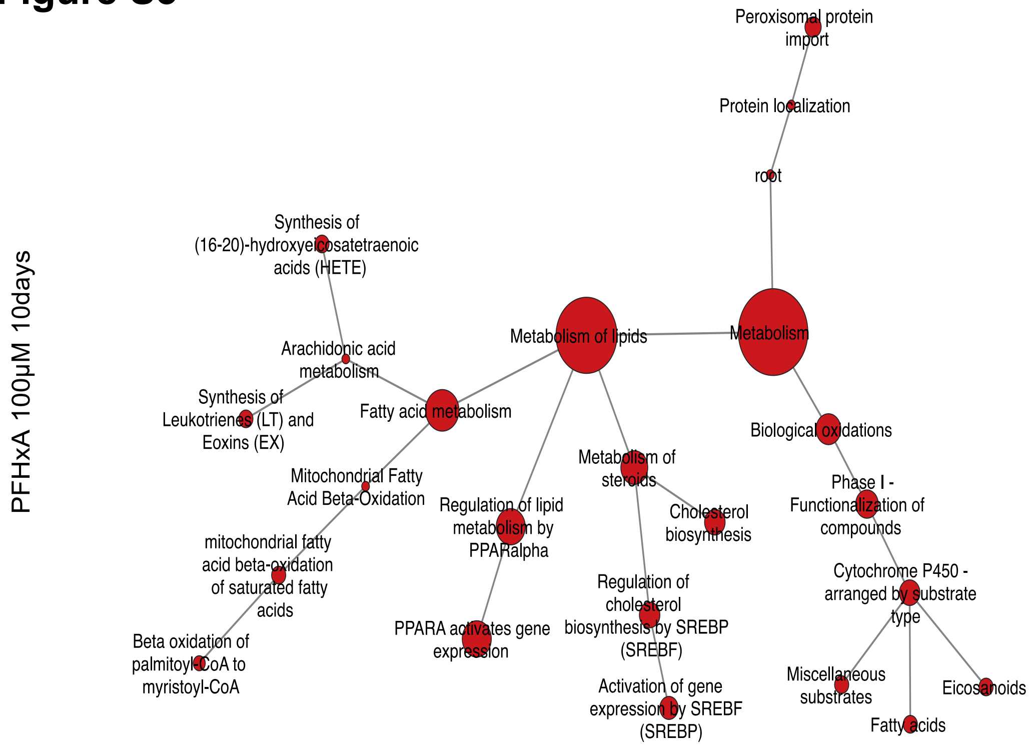

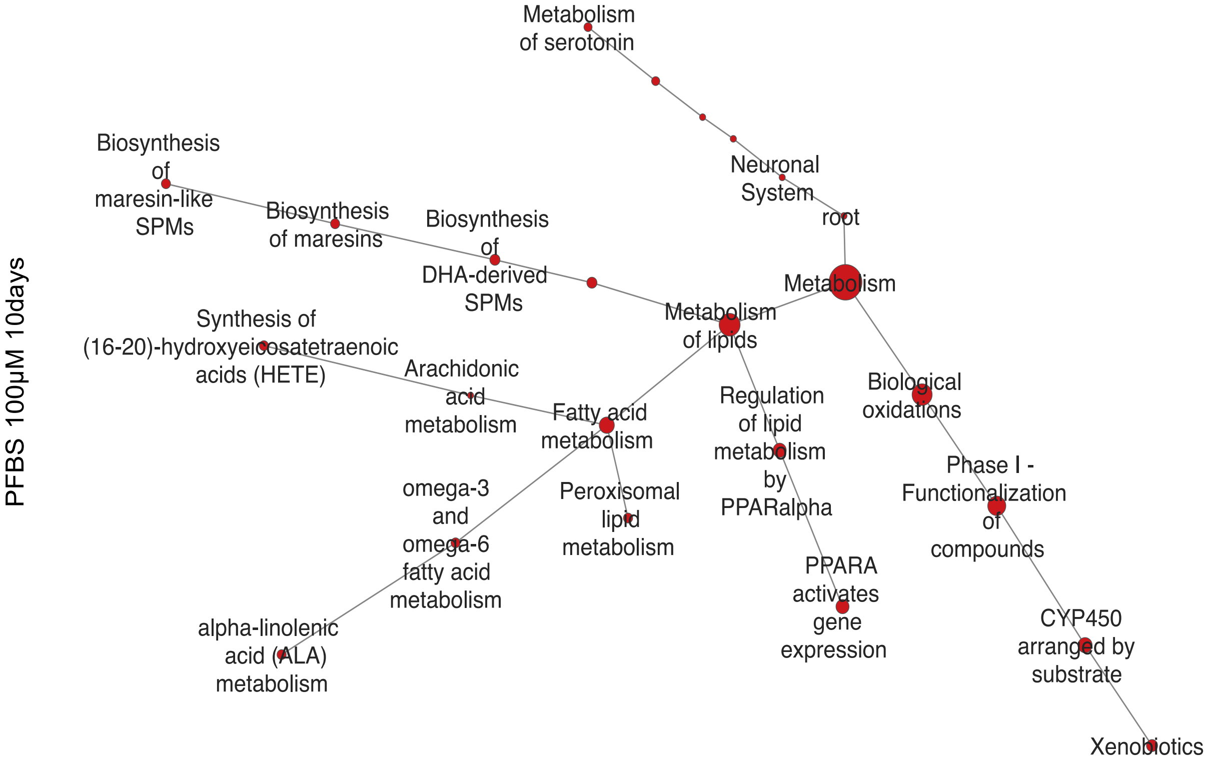

#### Figure S5

PFHpS 50µM 10days

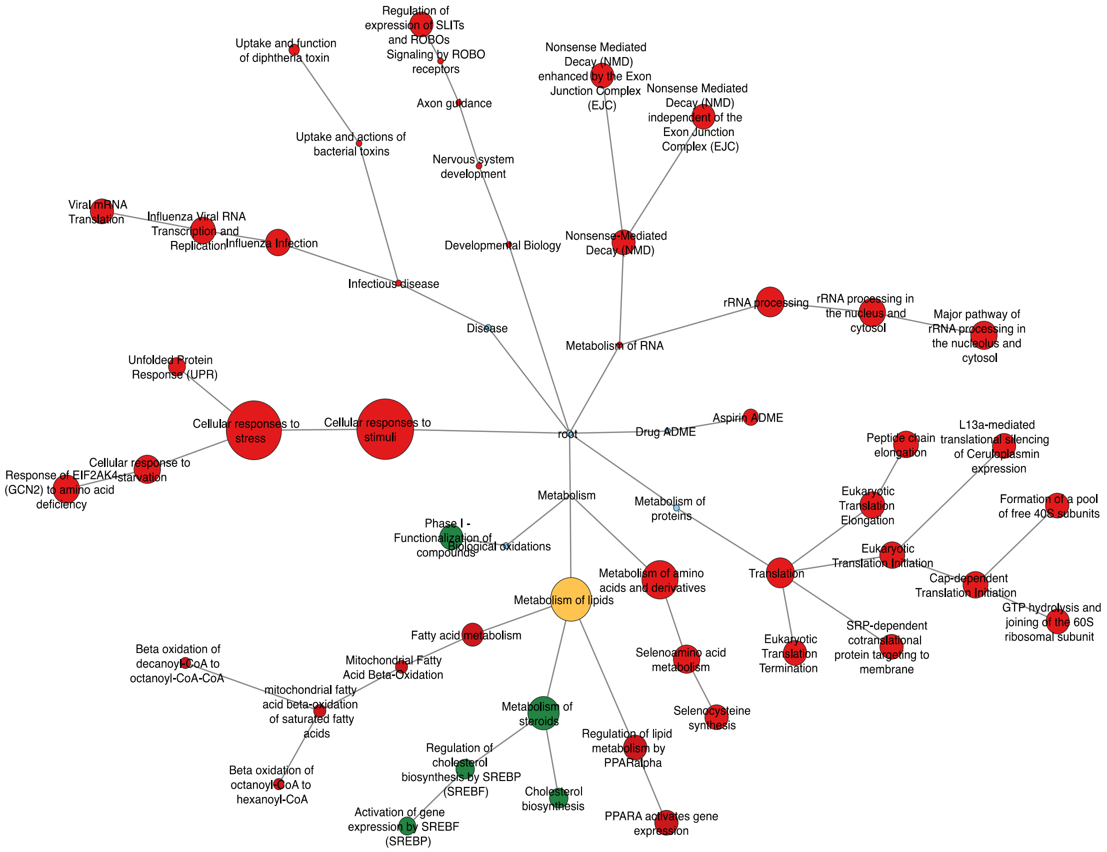

#### Figure S6

PFOA 20mg/kg/day 5days

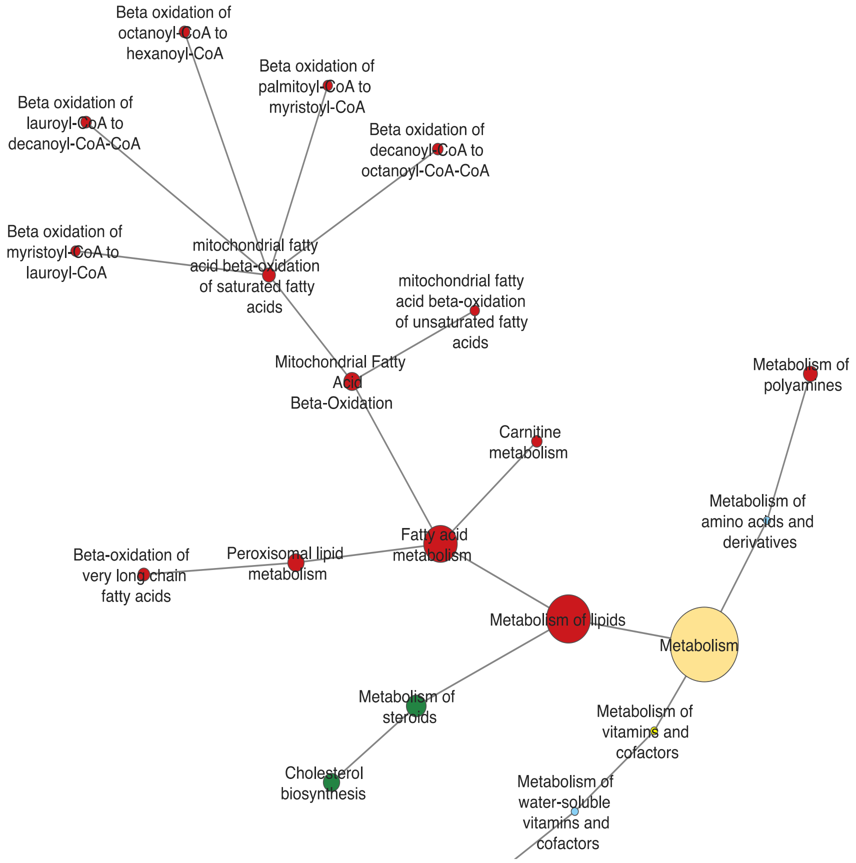
